# Integrative analysis of Hydra head regeneration reveals activation of distal enhancer-like elements

**DOI:** 10.1101/544049

**Authors:** Rabi Murad, Aide Macias-Muñoz, Ashley Wong, Xinyi Ma, Ali Mortazavi

## Abstract

The cnidarian model organism Hydra has long been studied for its remarkable ability to regenerate its head, which is controlled by a head organizer located near the hypostome. Cnidarians and bilaterians diverged about 600 millions years ago but the gene contents of species of both phyla are surprisingly similar despite divergent morphologies and functions. While little is known about the role of *cis*-regulatory elements in cnidarians, understanding gene regulatory mechanisms in cnidarians can potentially shed light on metazoan evolution. The canonical Wnt pathway plays a central role in head organizer function during regeneration and during bud formation, which is the asexual mode of reproduction in Hydra. However, it is unclear how shared the developmental programs of head organizer genesis are in budding and regeneration. Time-series analysis of gene expression changes during head regeneration and budding revealed a set of 298 differentially expressed genes during the 48-hour head regeneration and 72-hour budding time-courses. In order to understand the regulatory elements controlling hydra head regeneration, we first identified 27,137 open-chromatin elements that are open in one or more sections of organism. We used histone modification ChIP-seq to identify 9998 candidate proximal promoter and 3018 candidate enhancer-like regions respectively. We show that a subset of these regulatory elements is dynamically remodeled during head regeneration and identify a set of transcription factor motifs that are enriched in the enhancer regions activated during head regeneration. Our results show that Hydra displays complex gene regulatory structures of developmentally dynamic enhancers, which suggests that the evolution of complex developmental enhancers predates the split of cnidarians and bilaterians.

## Introduction

Hydra belongs to the phylum *Cnidaria* that consists of ∼10,000 species divided into two major groups: Anthozoa (comprising of sea anemones, corals, and sea pens) and Medusozoa (sea wasps, jellyfish, and Hydra). The hallmark traits of cnidarians are their external radial symmetry and the nematocyte, stinging cells used for predation. Unlike the more common bilaterians such as vertebrates and insects with left-right symmetry, cnidarians consist of two germ layers (endoderm and ectoderm) and have a single body axis called the oral-aboral axis. An adult Hydra has a simple structure consisting of a cylindrical tube with an apical head and a basal foot. The epithelial cells of both the ectoderm and endoderm of the body column are constantly in the mitotic cycle (1). As a consequence, tissue is continuously displaced towards and sloughed off at the two extremities (1). To maintain the structure of an adult hydra in this context, a small set of cells referred to as the head organizer are located in the hypostome in the upper part of the head (2–4). The head organizer actively maintains the pattern and morphology of the animal in the context of its tissue dynamics by signaling neighboring cells to adopt differentiated states appropriate to the head (hypostome and tentacles). When a hydra is bisected anywhere along the top 1/3 of the body column, a head regenerates at the apical end of the lower part of the bisected animal that first involves head organizer formation (5,6).

The Hydra *Wnt* and *TCF* genes of the canonical Wnt pathway are expressed in the hypostome where the organizer is located (7). A critical component of organizer formation is *β-catenin* (8). When Hydra are treated with alsterpaullone, which blocks the degradation of *β-catenin* by *GSK3β* (9), the level of *β-catenin* is elevated throughout the body column and results in numerous head organizers forming all along the body column (10). In addition, a number of other genes have been shown to affect, or be associated with head organizer formation. These include *Goosecoid* (11), *Brachyury* (12), *Forkhead/HNF-3b* (13), and *Chordin* (14).

The head organizer also arises during bud formation, which is Hydra’s asexual form of reproduction. Under normal physiological conditions Hydra reproduce asexually through budding in the lower body column area. During the initial stage of bud formation, a head organizer is formed in the budding zone, and subsequently directs the formation of a bud, which eventually develops into an adult Hydra. In addition to the their role in the formation and maintenance of the head organizer at the hypostome, Hydra Wnt genes are also involved in the budding process as they are expressed at the budding zone where the presumptive bud arises and in the hypostome of the growing bud (7). Since both regeneration and budding involve the formation of a head organizer, a natural question is the extent to which the two gene expression programs are similar. Specifically, what are the common and regeneration-specific (or budding-specific) sets of genes involved in head organizer genesis during head regeneration and budding, and subsequently its activity and maintenance? RNA-seq has enabled gene expression profiling, full-transcript assembly, allele-specific expression profiling and RNA-editing studies (15). During the last 5 years, RNA-seq has been used to assemble a transcriptome of Hydra (16), to characterize the transcriptome and proteome of Hydra during head regeneration (17), and to profile the small non-coding RNA repertoire of Hydra (18).

From a phylogenetic perspective cnidarians and bilaterians diverged ∼600 millions years ago (19). Therefore the study of cnidarians provides potential opportunities for elucidating key aspects of metazoan evolution such as the formation of mesoderm, bilaterian body plan and the nervous system. Considering the important evolutionary insights that can be obtained from comparison of cnidarians and bilaterians, genome sequencing and functional genomic studies of cnidarians have been at the forefront. Such efforts culminated in the sequencing of the genomes of the anthozoan *Nematostella vectensis* (20) in 2007 and of the medusozoan *Hydra vulgaris* (21) in 2010. A rather surprising finding of the genome sequencing of Hydra and *Nematostella* was that the gene contents of these basal metazoans are similar to those of bilaterians (20,21). This finding led to speculation that the difference in the body plans of cnidarians and bilaterians is due to differences in gene regulation (22) based on findings that body plan evolution is often a consequence of changes in gene regulation (23) and differences in *cis*-regulatory elements among even closely related species (24,25).

Gene expression at the transcriptional level is controlled by DNA sequences called *cis*-regulatory modules (CRMs). CRMs are short genomic regions that play specific roles in controlling the expression of their associated genes. CRMs act as binding sites for sequence-specific as well as general transcription factors (TFs) that mediate the expression of genes. CRMs are classified into promoters and enhancers. Promoters are short genomic regions overlapping the transcription start sites (TSSs) of genes and act as binding sites for general transcription factors that complex with RNA Polymerase II (26). Enhancers are *cis*-acting 200-1000 base pair (bp) DNA segments that contain multiple transcription factor binding sites (TFBSs) and control the expression of target genes (27). They are sometimes located tens to hundreds of kilobases (Kb) away from their associated genes (27) and can be located up to 1 megabase (Mb) from their targets (28). Once the right combination of transcription factors are bound to the TFBSs at an enhancer, the enhancer region is brought within close proximity of the target gene promoter by DNA looping (29).

A comparison of the gene regulatory landscapes of genomes in the cnidarians and bilaterians would require systematic genome-wide mapping of *cis*-regulatory elements within sequenced cnidarian genomes. So far, this has only been attempted in *Nematostella*, leading to identification of over 5000 enhancers and the surprising finding that the gene regulatory landscape of *Nematostella* is at least as complex than those of bilaterians (22). Although studies in mammalian model systems show that enhancers evolve at a more rapid pace than the coding sequences, it remains to be determined if enhancer evolution is an important agent of metazoan evolution over long evolutionary periods.

The first step in a comparative study of enhancers in cnidarians is a genome-wide identification of enhancers and other regulatory elements. The most reliable approach involves experimental methods to identify genomic regions containing features associated with enhancers (30). Experimental methods for mapping promoters and enhancers are based on detecting specific histone modifications that are associated with each type CRM type as well as increased accessibility of such regions due to localized depletion of nucleosomes. Specific combinations of posttranslational modifications of histone proteins are associated with either promoters or enhancers. For example, histone H3 in promoter regions are associated with high levels of trimethylation or dimethylation at Lysine 4 (H3K4me3 and H3K4me2, respectively) in fungi, plants, and animals while high levels of H3K4me2 and H3K27ac (H3 Lysine 27 acetylation) but low levels of H3K4me3 are found at active enhancer regions in bilaterians. Chromatin immunoprecipitation followed by sequencing (ChIP-seq) using antibodies that recognize specific histone modifications can be used to map the locations of candidate active promoters and enhancers genome-wide. These candidate promoters and enhancers are furthermore located in genomic regions relatively depleted of nucleosomes, which allows transcription factors the ability to bind their binding site. These regions are more readily digested enzymatically due to easy accessibility compared to tightly wound “closed” chromatin. This is the basis for high-throughput assays, such as DNase-seq (31–33) and ATAC-seq (34), which preferentially digest open-chromatin regions with the enzymes DNase and Tn5 transposase respectively when treated for a short period of time. In bilaterians, changes in open chromatin accessibility are observed during differentiation and development.

In this study we used RNA-seq to characterize genome-wide gene expression patterns during Hydra head regeneration and budding. In addition, we profiled the open-chromatin elements of Hydra using ATAC-seq during a 48-hour time-course of head regeneration and body map as well as generated corresponding ChIP-seq experiments of three histone modifications (H3K4me2, H3K4me3, and H3K27ac) to map genome-wide candidate promoter and enhancer-like elements in Hydra (Fig 1). We analyzed the resulting differentially expressed genes to assess the common and divergent sets of genes between head regeneration and budding in Hydra as well gene signatures of Hydra body tissues such as hypostome. The integrative analysis of ATAC-seq and ChIP-seq datasets allowed us to predict 9998 candidate promoters and 3018 candidate enhancer-like elements in the Hydra genome. We find evidence for extensive chromatin remodeling of the regenerating head tissue and we identify a set of motifs for specific transcription factors that are enriched in the enhancer regions that are activated during remodeling.

**Figure 1:**
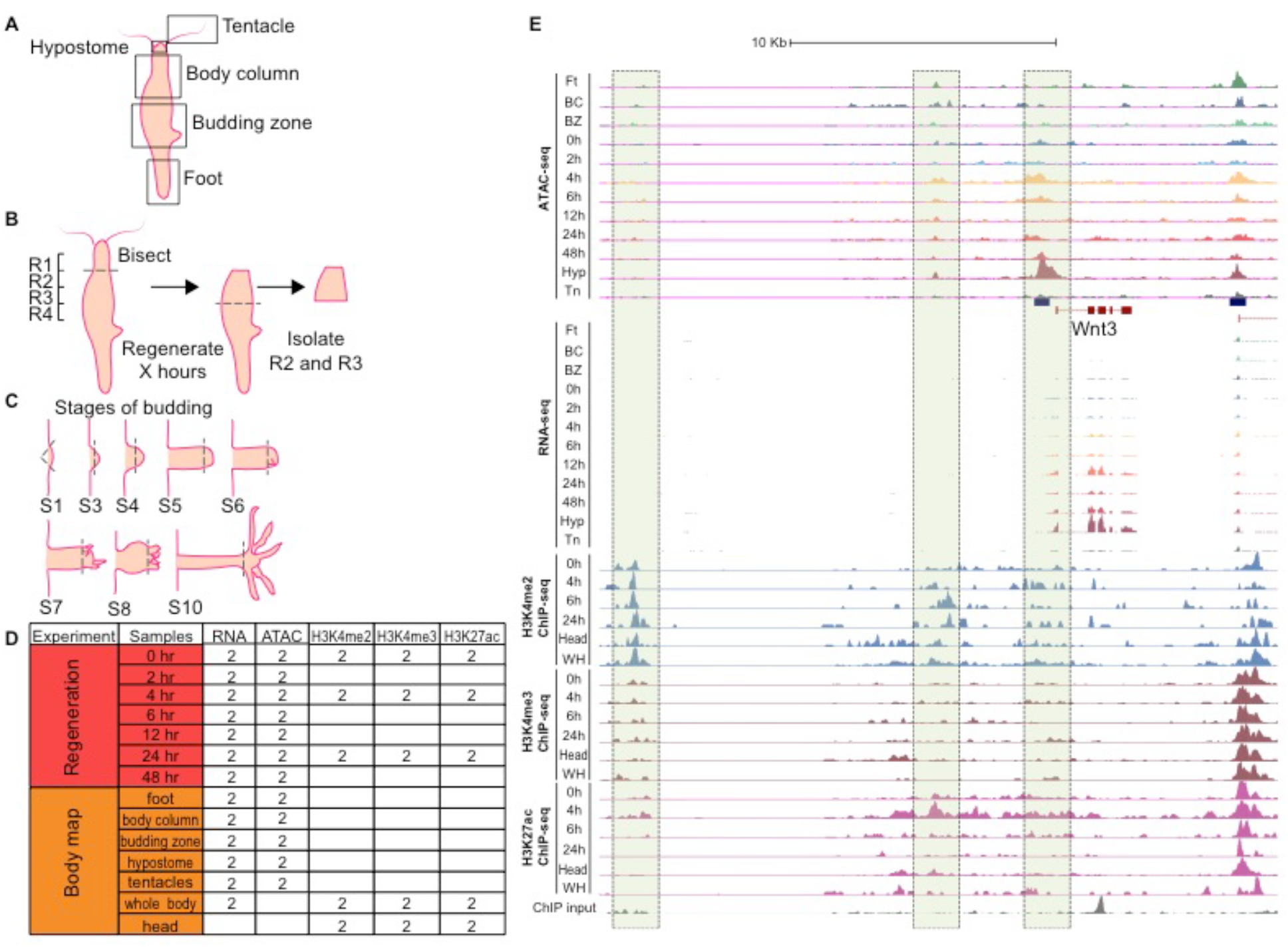
Outline of RNA-seq, ATAC-seq, and ChIP-seq experiments. **(A)** Five Hydra body parts were isolated for RNA-seq and ATAC-seq to create a body map. **(B)** Hydra heads were bisected at the boundary of regions R1 and R2 and allowed to regenerate for specific time periods (0, 2, 4, 6, 12, 24, and 48 hours). The regions R2 and R3 were isolated for RNA-seq to measure gene expression and ATAC-seq to map chromatin accessibility. **(C)** Hydra bud heads at various stages of budding (S1, S3, S4, S5, S6, S7, S8, and S10) were bisected and total RNA was extracted for RNA-seq. **(D)** Number of Hydra biological replicate samples assayed by RNA-seq, ATAC-seq, and ChIP-seq. Two biological replicates were also obtained for each stage of budding. **(E)** Genome browser signal tracks for RNA-seq, ATAC-seq and ChIP-seq data for the Wnt3 locus. The Wnt3 promoter and an upstream “enhancer” region gain hypersensitivity along the head regeneration time course as gene expression is turned on.

## Results

### Hydra head regeneration and budding time courses and body map

We performed time-course RNA-seq, ATAC-seq, and ChIP-seq experiments in Hydra during head regeneration and budding as well as certain body parts. To generate a “body” map for Hydra, body column, budding zone, foot, tentacle, and hypostome tissues were collected for RNA-seq and ATAC-seq while whole animal and head tissues were collected for ChIP-seq (Fig 1A, 1D). For head regeneration, animals were bisected between the regions R1 and R2 and allowed to regenerate for certain time periods (0, 2, 4, 6, 12, 24, and 48 hours) and the regenerating tips were collected for RNA-seq, ATAC-seq, and ChIP-seq (Fig 1B, 1D). For budding, the heads of developing buds at specific stages of budding (S1, S3, S4, S5, S6, S7, S8, and S10) as classified by Otto *et al.* (35) were collected for RNA sequencing (Fig 1C, 1D). All experimental samples were done in two biological replicates for reproducibility. The sample libraries were built using the same protocol and the datasets were processed uniformly.

### Hydra body map transcriptome

A principal component analysis (PCA) of RNA-seq samples from tentacles, hypostome, body column, budding zone, and foot showed that each of the libraries groups by body part (Fig 2A). The foot, tentacle and hypostome were more distinct from each other than the body column and budding zone, which lie near each other on the PCA plot (Fig 2A). We found 217 genes differentially expressed (DE) between the body column and budding zone, 1146 between the body column and foot, 847 between the foot and budding zone, 4244 between the budding zone and the tentacles, 1836 between the foot and hypostome, 1774 between the body column and hypostome, 4478 between the body column and tentacles, 1887 between the budding zone and the hypostome, 3421 between the foot and tentacles, and 2760 between the hypostome and tentacles (Table S1). 204 genes were uniquely upregulated in the hypostome, 837 in the tentacles, 58 in the body column, one in the budding zone, and 259 in the foot (Table S1). The low number of genes uniquely upregulated in the body column and budding zone is likely due to the similarity in expression profiles for the two tissues.

**Figure 2:**
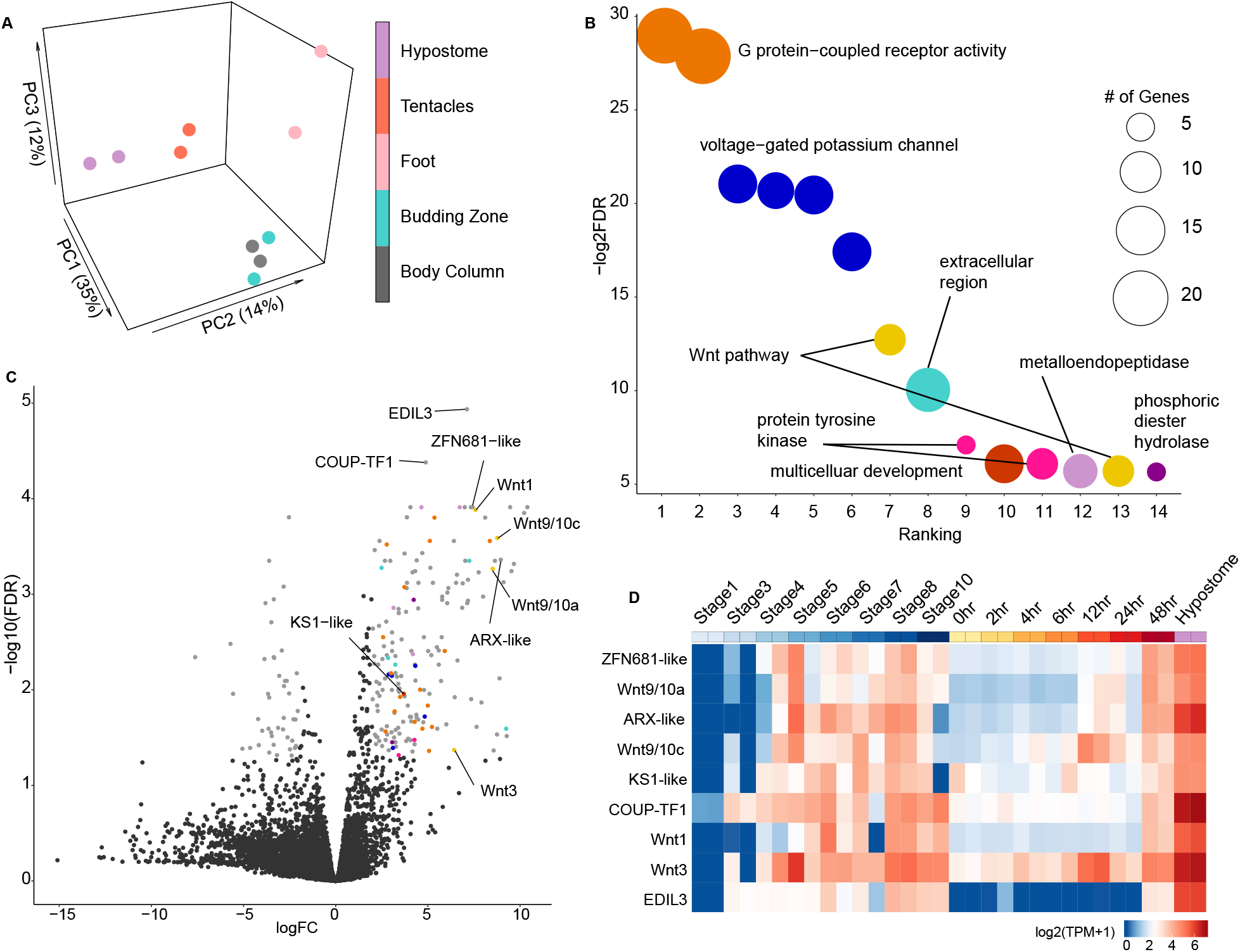
Gene expression of hypostome specific genes. **(A)** PCA plot showing the distinct expression profiles of hypostome, tentacles, body column, budding zone, and foot gene expression. **(B)** Functional enrichment analysis of genes upregulated in the hypostome using BLAST2GO and an FDR>0.05. Circle size represents number of genes and color represents similar functions. **(C)** Volcano plot comparing hypostome versus other tissues, colors represent significance (light grey) and functions (similar to those in 2B). **(D)** Heatmap showing changes in gene expression of key developmental genes from panel C upregulated in the hypostome throughout stages of budding (stage1-10) and time points in regeneration (0hr-48hr).

Genes uniquely upregulated in the hypostome were enriched for functions in G-protein coupled receptor activity, voltage-gated potassium channel activity, Wnt signaling pathway, extracellular region, protein tyrosine kinase activity, multicellular organism development, metalloendopeptidase activity, and phosphoric diester hydrolase activity (Fig 2B, Table S2). Genes enriched for functions in Wnt signaling and multicellular organism development included *Wnt3, Wnt1, Wnt9/10C*, and *KS1-like* (Table S3). Other potential genes of interest that were highly expressed in the hypostome relative to other tissues (low False Discovery Rates (FDR) and high log fold change (logFC)) were *EGF-like repeat and discoidin I-like domain-containing 3, COUP-TF1, Homeobox ARX*, and *zinc finger 681-like* (Fig 2C).

We expected to detect Wnt genes upregulated in the hypostome since Wnt genes have been implicated in the Hydra head organizer (36). While much attention has been given to *Wnt3* due to its role in head patterning and hypostome regeneration, six additional *Wnt*s (*1, 7, 9/10a, 9/10c, 11*, and *16*) are also expressed in the adult hypostome and the head organizer in budding and regeneration. In our investigation we found *Wnt3, Wnt1, Wnt9/10a*, and *Wnt9/10c* upregulated in the hypostome relative to other tissues in the adult Hydra. The reason that we did not detect the remainder of the Wnt genes as Hypostome-specific is because *Wnt11* is expressed in all tissues and *Wnt8* and *Wnt5a* are highly expressed in both the hypostome and the tentacles. We inspected the expression dynamics of these genes during budding and regeneration (Fig 2D). In a previous study that used *in situs, Wnt3* began expression at 1.5 hours after decapitation and was expressed at all stages of budding (36). We observed that *Wnt3* expression was present at low levels at 0h after decapitation and increases with time reaching high levels at 12h and 48h (Fig 2D). Similarly, for budding, expression increases throughout stages of budding with little to no expression at stage 1 (Fig 2D). *In situs* showed that *Wnt1* expression appeared at 3h but did not reach high levels until 6h during regeneration; *Wnt1* was also expressed at all stages in budding (36). Using RNA-seq, we noticed different patterns of *Wnt1* expression. During regeneration, *Wnt1* had low expression until 48h and during budding expression was low until stage 5 (Fig 2D). Lastly, *in situs* showed the *Wnt9/10a* and *Wnt9/10c* began expression at 6h and increased in expression at 12h post bisection; *Wnt9/10a* and *c* were also expressed at all staged of budding (36). RNA-seq showed that expression of *Wnt9/10a* and *Wnt9/10c* increased during regeneration. *Wnt9/10a* only reached high expression at 48h while *Wnt9/10c* had high expression at 12h (Fig 2D). During budding, *Wnt9/10a* and *Wnt9/10c* expression began increasing at stage 3 and 4 and reached high expression at stage 8, after which expression seemed to decrease (Fig 2D). While *in situs* help us localize where genes are expressed, these assays are absolute and can be sensitive to lowly expressed genes. Our results vary from previous *in situ* studies because we are able to quantify gene expression during different stages of budding and regeneration. Through these methods, we can better detect the dynamics of *Wnt* signaling to further understand head regeneration in hydra.

*Ks1* is another gene with a role in hydra head formation (37). During head regeneration, *Ks1* begins to be expressed 2 days after decapitation and is highly expressed at 4 days in the adult Hydra (37). In our study, we identified a *Ks1-like* gene highly expressed in the hypostome relative to other tissues (Fig 2C). We found expression of *Ks1-like* to be somewhat consistent during regeneration with expression increasing at 48h (Fig 2D). In addition, *Ks1-like* expression began to increase at stage 4 of budding and was highly expressed during hydra bud stage 6, 7, and 8. Compared to the Wnt genes, *Ks1-like* has a more similar trajectory in regeneration to *Wnt3* but increased at a much slower rate. Based on the expression patterns that we observe during regeneration and budding, *Ks1-like* expression follows that of *Wnt3* suggesting these genes may exist in a similar network. Continued high expression in the hypostome of the adult Hydra also suggests that *Ks1-like* is actively transcribed and may play a role in the head organizer.

Additional interesting genes in the hydra hypostome are those involved in transcription such as *EGF-like repeat and discoidin I-like domain-containing 3 (EDIL3), COUP-TF1, Homeobox aristaless-like* (*ARX-like*), and *zinc finger 681-like* (*ZFN681-like*). We focus on these genes because they have a high log fold change and high FDR. The functions of *EDIL3* and *ZFN681* are not known in Hydra but zinc finger proteins typically function in binding RNA or DNA and stabilizing protein-protein interactions. *EDIL3*, in humans, functions as an enhancer and encodes an integrin protein. During regeneration, *ZFN681-like* had expression similar to *Wnt1*, low expression throughout 0-24h and increased expression at 48h (Fig 2D). Conversely, EDIL3 has no expression in the first 24h and expression began at 48h (Fig 2D). Another gene highly expressed in the hypostome is *COUP-TF1* which has implications to function in neurogenesis. A *COUP-TF* gene family member was found expressed in cells that lead to nematocytes and neurons (38). Due to the overexpression of *COUP-TF1* in our study, this gene may function in neurons that are high in number or condensed in the hypostome (Fig 2C). Expression of *COUP-TF1* increased throughout regeneration and budding. During regeneration, expression is low until 48h and during budding average expression is similar at stage 5 to later stages (Fig 2D). Lastly, we found an aristaless-like gene highly expressed in the hypostome whose expression during budding was similar to that of *Wnt3* and similar to *Wnt9/10a* during regeneration (Fig 2D). The particular function of this homeobox gene has yet to be described but a member of the aristaless gene family functions in tentacle formation in hydra (39). The expression of *Arx-like* in our data set suggests this homeobox gene has a role in head patterning.

### Time-series analysis of Hydra head regeneration and budding transcriptomes

Hydra head regeneration is a classic example of the critical role of signaling pathways in axial patterning and head organizer formation and maintenance (6). Head regeneration has been studied using both developmental approaches (7) and more recently using genomic approaches (17). However, less is known about the extent to which the regeneration and budding gene regulatory programs overlap. We used PCA to compare globally the regeneration and budding transcriptomes as well as certain body map tissues. Tentacles cluster separate from the regeneration time points, budding stages, body column, and hypostome samples along principal component 1 (PC1) which accounts for the highest amount of variance (26%) (Fig 3A). Such sharp clustering and separation of regeneration and budding samples from tentacles along PC1 indicates that there is a large set of genes whose expression are very specific only to tentacles. In contrast, principal components 2 and 3 (PC2 and PC3) reveal that the time courses of head regeneration and budding follow different trajectories to converge at a single point corresponding to the hypostome samples (Fig 3A).

**Figure 3:**
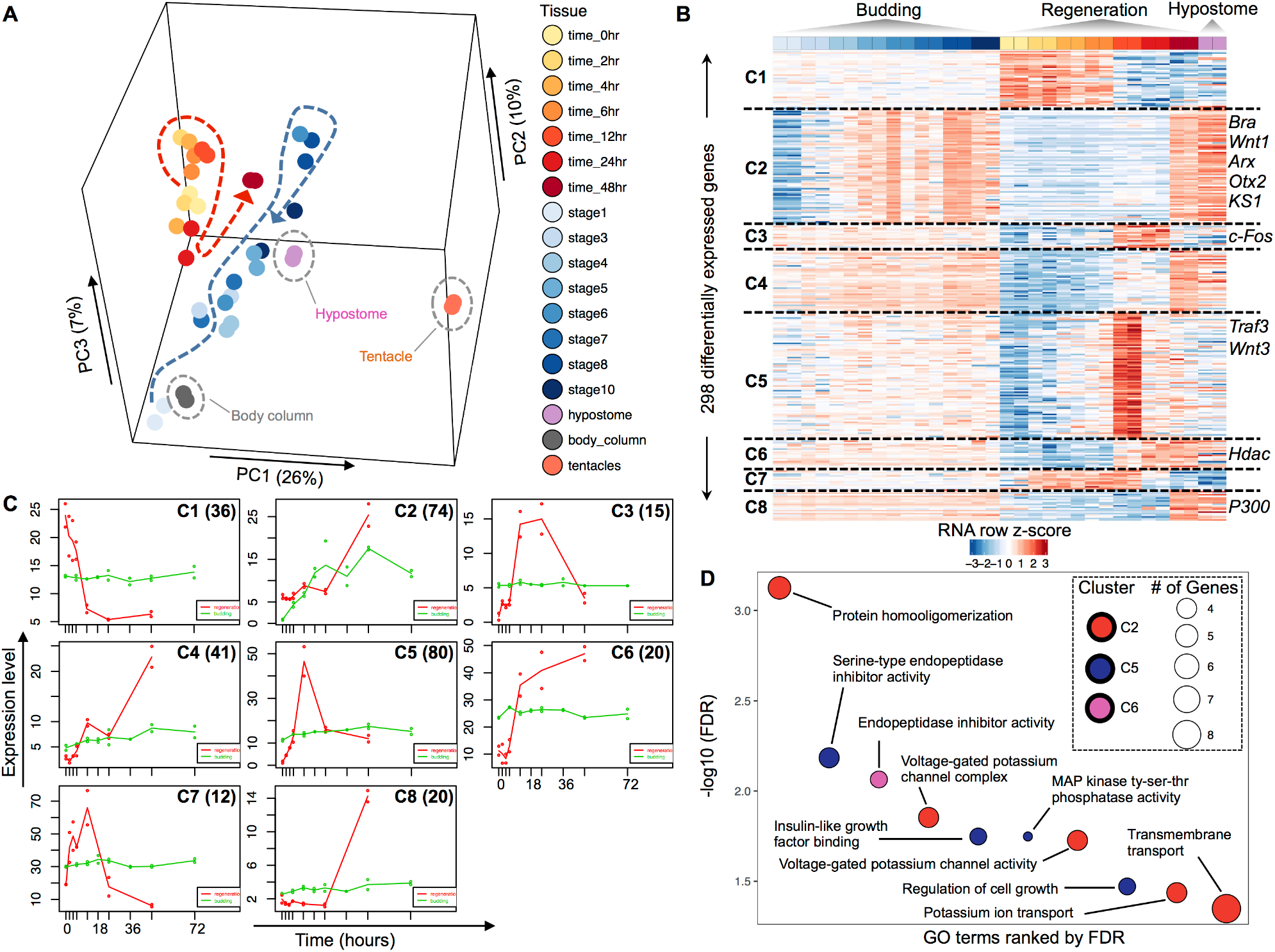
Comparative analysis of gene expression between head regeneration and budding in Hydra. **(A)** PCA of Hydra gene expression during head regeneration, budding, and in selected body parts showing the developmental trajectories of head regeneration and budding. **(B)** Heatmap of 298 differentially expressed genes, grouped in 8 clusters based on similar expression dynamics during head regeneration and budding. **(C)** Median expression profiles of clusters in the time courses of regeneration (red profiles) and budding (green profiles). The clusters correspond to panel B and the number of genes in each cluster are shown inset. **(D)** Representative enriched GO terms for the clusters of differentially expressed genes in Fig. 3B. Significance level is at FDR of 0.05. Clusters 1, 3-4, and 7-8 did not have any enriched GO terms.

The PCA analysis of Hydra head regeneration and budding shows that there are sets of genes specific or common to both regeneration and budding time-courses. For a higher resolution analysis to determine these gene sets, we performed a time-series analysis of the regeneration and budding transcriptomes using maSigPro (40) to find clusters of differentially expressed genes that have similar expression profiles. This analysis identified 298 DE genes that form eight non-redundant clusters (Fig 3B). DE genes in seven clusters (C1, C2-C8) are expressed at a steady state during the budding time-course but show complex dynamic temporal changes during regeneration (Fig 3B). Genes in cluster C2 show increasing expression along both time courses (Fig 3B). Median expression profiles of DE genes in each clusters are shown in Fig 3C. Some of the hypostome marker genes such as *Wnt1, Wnt3, ARX*, and *KS1* are upregulated during different stages of regeneration (Fig 3B, 2C).

The genes in each of the eight distinct clusters (Fig 3B) were tested for enrichment of gene ontology (GO) terms using Blast2GO’s Fisher exact test (FDR ≤ 0.05). Genes in three clusters (C2, C5, and C6) were enriched for GO terms while five clusters (C1, C3, C4, C7, and C8) did not have any significantly enriched GO terms (Fig 3D). Cluster 2, consisting of 72 genes upregulated in mid-to-late stages of both head regeneration and budding, had enriched GO terms related to “protein homooligomerization”, “voltage-gated potassium channel complex”, “voltage-gated potassium channel activity”, “potassium ion transport”, and “transmembrane transport”. Cluster 5 (80 genes) was upregulated during early head regeneration time points (4-12 hours post-bisection) and was enriched in GO term such as “serine-type endopeptidase inhibitor activity”, “insulin-like growth factor binding”, “MAP kinase tyrosine-serine-threonine phosphatase activity”, and “regulation of cell growth”. Cluster 6 (20 genes) which increased in expression during regeneration was enriched in “endopeptidase inhibitor activity”.

### Mapping the open-chromatin landscape of Hydra

We sequenced the ATAC-seq libraries to an average depth of 20 million reads and mapped the reads onto the latest release of the Hydra genome (Hydra 2.0 Genome Project) in order to identify open-chromatin “peaks” in each dataset using Homer (41). Peak calls from each biological replicate were compared and only those that overlapped were retained for downstream analysis. The sets of peaks from all samples in the regeneration time-course and body-map were merged to obtain a consolidated set of 27,137 peaks.

We classified the 27,137 open-chromatin elements according to their genomic locations with respect to the annotated genes. Since no prior knowledge about the locations of regulatory elements in Hydra was available, we defined four classes of open-chromatin elements as follows: proximal promoter (peaks within ±2 Kb of transcript start sites), intergenic (peaks located between genes), intronic (peaks overlapping annotated introns), and exonic (peaks overlapping annotated exons) (Fig 4A). Using this classification scheme, we identified 9998 proximal promoter open elements, 8962 intergenic open elements, 6454 intronic open elements, and 1723 exonic open elements (Fig 4B). We next looked at the distribution of ATAC-seq signal in the four types of open-chromatin elements defined above (Fig S1). The proximal promoter open elements possess the highest amount of signal followed by the intergenic open elements. Most of the signal is located at the centers of the four types of peaks. Based on the genomic locations of the peaks and the enrichment of ATAC-seq signal at them and their distance from the annotated TSS, our set of open-chromatin elements provides candidate promoter (near TSS) and enhancer-like elements (intergenic). However, we cannot exclude that some intergenic elements represent unannotated TSSs without using additional evidence such as the ratio of H3K4me3 to H3K4me2 from the ChIP signal.

**Figure 4:**
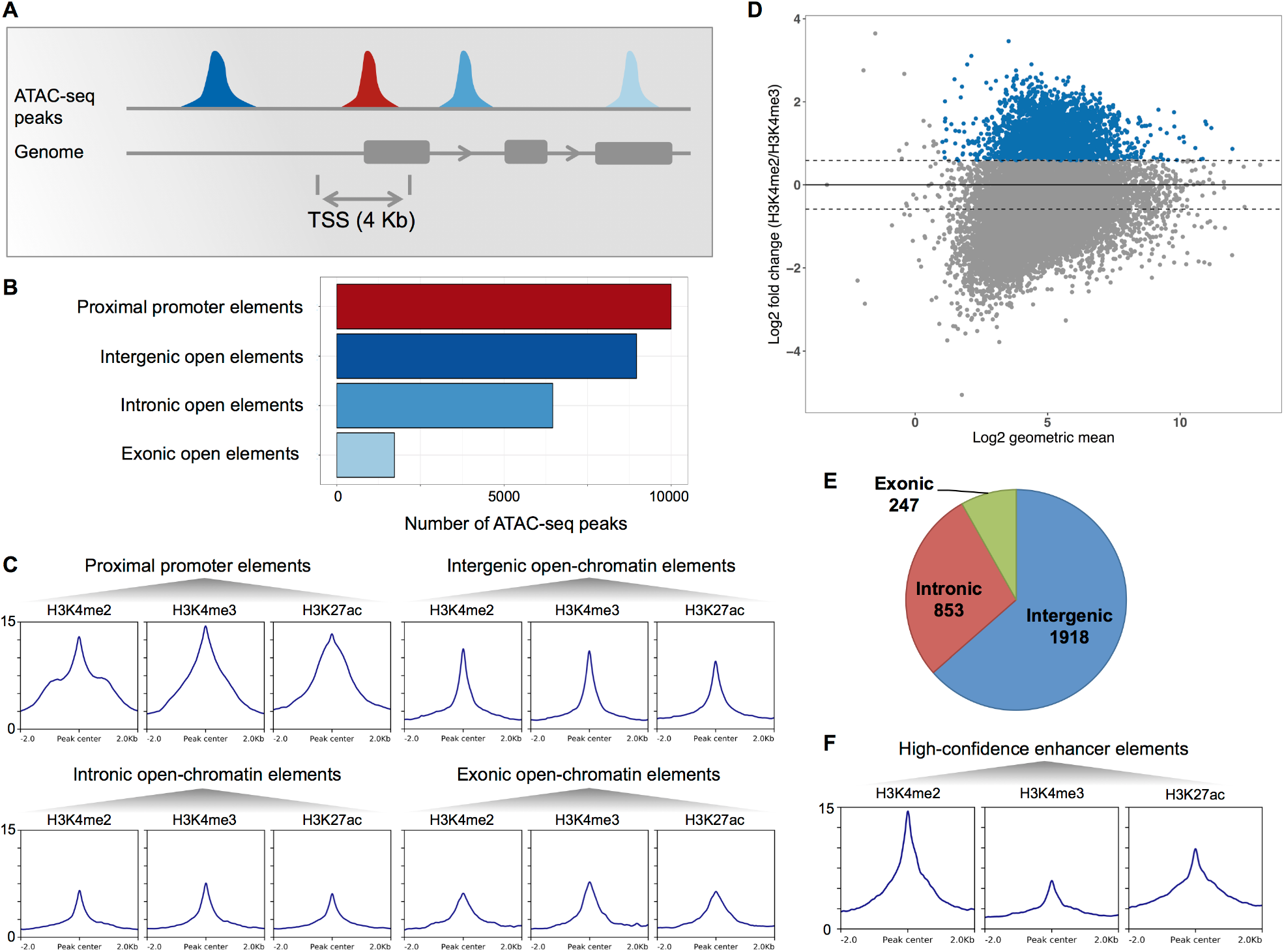
Identification of a high-confidence set of Hydra regulatory regions using ATAC-seq and ChIP-seq data. **(A)** Distribution of open-chromatin elements based on their genomic locations. **(B)** The 27,137 replicated ATAC-seq peaks found in one or more replicated samples were classified based on overlap with gene loci. The proximal promoter regions were defined as regions within 2 Kb of start of genes. **(C)** Enrichment of H3K4me2, H3K4me3, and H3K27ac histone modifications normalized signals at each type of open-chromatin elements. **(D)** 3018 open-chromatin elements (blue) elements with at least 50% higher H3K4me2 than H3K4me3 signal form a set of high-confidence candidate enhancer-like. **(E)** Genomic distribution of candidate enhancer-like elements. **(F)** Enrichment of H3K4me2, H3K4me3, and H3K27ac normalized signals at the candidate enhancer-like elements.

### Classification of open-chromatin elements using histone modifications

Using our ATAC-seq datasets, we obtained a set of 27,137 open-chromatin elements that were classified into four groups based on their genomic locations (Fig 4B). We used ChIP-seq (42) in several of the corresponding time points of the head regeneration time-course and body parts as well as whole animal (Fig 1D) to generate histone modification profiles. We used antibodies that detect H3 dimethylation of Lys 4 (H3K4me2), trimethylation of Lys 4 (H3K4me3), and acetylation of Lys 27 (H3K27ac). The histone modifications H3K4me3 and H3K4me2 are known to mark chromatin at the promoter regions with a higher ratio of H3K4me3 to H3K4me2, whereas high H3K4me2 with low H3K4me3 predominantly mark the enhancer regions and H3K27ac marks active regulatory regions (43).

We computed and compared the normalized enrichment of the H3K4me2, H3K4me3, and H3K27ac at the peaks from the four sets of open-chromatin elements classified based on their genomic locations (Fig 4C). The proximal promoter open-chromatin elements showed the highest enrichment of H3K4me2, H3K4me3, and H3K27ac with a slightly higher enrichment of H3K4me3 (Fig 4C). Thus, the chromatin marks provide further evidence for the proximal promoter open-chromatin elements as candidate promoter regions in the Hydra genome. We expected higher enrichment of H3K4me2 compared to H3K4me3 at the remaining three classes of open-chromatin elements (intergenic, intronic, and exonic) since these were non-TSS overlapping. We observed almost equal (intergenic, Fig 4C) or slightly lower enrichment of H3K4me2 (intronic and exonic, Fig 4C) at these regions. A reason for the discrepancy in the relative enrichments of H3K4me2 and H3K4me3 at the non-TSS open-chromatin element sets could be the inclusion of peaks overlapping the non-annotated TSS regions. Therefore, we used the relative enrichment of H3K4me2 over H3K4me3 to score the peaks and identify candidate enhancer regions (Fig 4D). We defined H3K4me2 enriched peaks as having minimum 50% or higher enrichment of H3K4me2 signal over H3K4me3 (Fig 4D). This strategy led to the identification of 3018 ATAC-seq peaks which are predominantly intergenic (1918/3018) followed by intronic (853/3018) and exonic open-chromatin elements (247/3018) that have higher H3K4me2 than H3K4me3 signal (Fig 4E). Comparison of the histone mark signals at the 3018 peaks reveals considerable enrichment of H3K4me2 relative to the H3K4me3 mark (Fig 3F). Therefore, the set of 3018 open-chromatin elements, based on their genomic locations and the enrichment of histone modifications, form the likeliest candidates for enhancer-like regions in the Hydra genome.

### Dynamics of open-chromatin elements during Hydra head regeneration

Hydra head regeneration is a dynamic process involving changes in expression of multiple genes related to Wnt signaling pathway (36), MAPK pathway (44), and response to injury (17) to name a few. An important question that remains unanswered is: How extensive is the remodeling of chromatin in Hydra genome in response to bisection and regeneration of a head? With a genome-wide set of open-chromatin elements obtained from data generated in this study, we explored the above question.

Dynamic remodeling of the chromatin during Hydra head regeneration time-course can be observed at the *Wnt3* gene locus (Fig 1E) which is known to be one of the earliest Wnt ligands expressed at the regenerating head (36). Open-chromatin signals appear at the Wnt3 promoter and upstream candidate enhancer-like regions as early as 4 hours post bisection (Fig 1E). We extended this analysis to the complete set of 27,137 open-chromatin elements by looking for differentially accessible (DA) elements genome-wide. Differential analysis of all the peaks identified 2870 DA open-chromatin elements, at 5% FDR and minimum 2-fold change, that form eight groups with distinct dynamic patterns when clustered (Fig 5A). The clusters reveal sets of open-chromatin elements specific to certain tissues or head regeneration time-course. For example, cluster 1 consists of open-chromatin elements specific to the foot, budding zone, and body column tissues of Hydra, while cluster 3 and 8 consist of elements that lose or gain accessibility during head regeneration respectively (Fig 5A).

**Figure 5:**
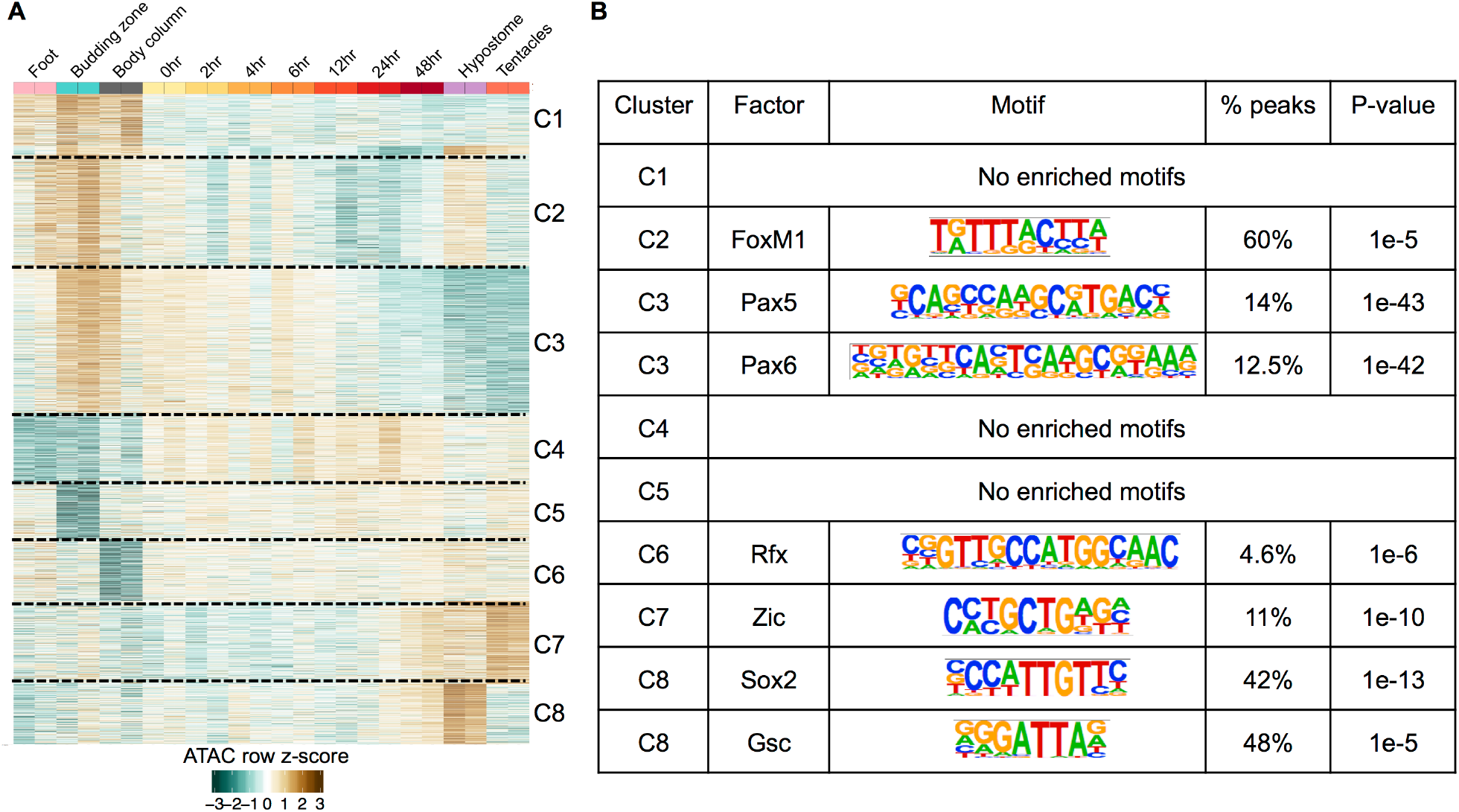
Dynamics and motif analysis of 2870 differentially accessible (DA) peaks. **(A)** Normalized reads per million (RPM) values for the 2870 DA peaks were converted to row z-scores and k-means clustered into 8 clusters based on the observed number of clusters when hierarchically clustered. **(B)** Transcription factor binding motifs enriched in open chromatin regions of each cluster in part A.

We next determined whether the tissue and regeneration time course specific clusters of the DA peaks were enriched for transcription factor binding sites. Five clusters of DA peaks (C2-C3, C6-C8) had enriched motifs for transcription factor binding sites while three clusters (C1, C4, and C5) had no enriched motifs. Open chromatin regions of cluster C2 (specific to foot, budding zone, body column, and hypostome) were enriched for binding site of FoxM1. Open chromatin regions specific to budding zone and body column that lose accessibility along regeneration time course (cluster C3) were enriched for the Pax5 and Pax6 binding motifs. Cluster 7, consisting of peak regions that gain accessibility in later stages of regeneration (24-48 hours post-bisection) and specific to hypostome and tentacles were enriched for Zic binding site which is known to play a role in specification of sensory nematocytes (45). Peak regions in cluster C8 that gain accessibility along the time course of regeneration were enriched for the binding sites of Sox2 and Goosecoid (Gsc). The Goosecoid homologue in Hydra is known to participate in head patterning (11). Overall, we found that the nearly ten percent of candidate regulatory elements that show dynamic changes during Hydra head regeneration are enriched in predicted transcription factor binding sites of developmental TFs.

## Discussion

We carried out time-course experiments in Hydra using RNA-seq, ATAC-seq, and ChIP-seq to compare gene expression during head regeneration and budding and to obtain a genome-wide view of open-chromatin landscape and remodeling in the Hydra genome in the context of head regeneration and body-map. To further classify the open-chromatin elements, we carried out ChIP-seq of three histone modifications (H3K4me2, H3K4me3, and H3K27ac) in a subset of these corresponding samples to annotate the open-chromatin elements as promoter-like or enhancer-like regions.

The time-course RNA-seq experiments in this study has shed light on genome-wide gene expression patterns during formation of the head organizer in Hydra during head regeneration and budding. The head organizer in Hydra is estimated to consist of 50-300 cells at the apical tip of the head. Single animal profiling at a greater temporal resolution should provide additional insights into the establishment of head organizer in different developmental scenarios in Hydra and the processes that initiate and maintain it. Whether the head organizer plays a role in sexual embryonic development is not known, although it is likely also involved in the development of the structure of the animal during embryogenesis. Future extensions of this study to the comparison with the head organizer formation during sexual embryogenesis will reveal the extent of reuse of the normal developmental program during head regeneration.

Identification of developmental genes such as *Wnt, Ks1* and *ARX* upregulated in the adult hypostome and their dynamic expression during regeneration and budding has implications for their role in the head organizer. As mentioned above, as Hydra grow and cells slough off, the head organizer must continuously be made anew through yet unknown signals and mechanisms. We predict that in adult Hydra, these head organizer “signals” must be expressed constantly for continuous cell differentiation. We also predict that these signals increase during head regeneration and budding. Interestingly enough, *Wnt1, ARX*, and *Ks1* have similar expression patterns that follow what we predict would happen as the head organizer is determined (Fig 3B). *Wnt3* is expressed in a different cluster due to coming on and peaking much earlier during regeneration and expression decreasing in the adult head (Fig 3B). These results imply that *Wnt1, ARX*, and *Ks1* maintain head organizer differentiation; they are highly expressed in the adult, expressed during most stages of budding and only in late stages of regeneration once cells have taken on a head organizer role. *Wnt3*, while also important to the head organizer and upregulated in the hypostome, is crucial during head regeneration. *Wnt3* is the first gene that comes on and may be triggering an increase of other Wnt and related genes. We propose that *Wnt3* is responsible for head organizer cell determination while other genes play a role in head organizer cell differentiation and maintenance.

In this study, we provide the first genome-wide atlas and analysis of open-chromatin elements in a cnidarian genome in the context of a developmental process. We can not determine which of our datasets (ATAC-seq or histone mark ChIP-seq) is more predictive of enhancers due to lack of a gold standard enhancer set in Hydra that could be used to evaluate our datasets. In mouse, chromatin accessibility (such as DNase-seq and ATAC-seq) and H3K27ac ChIP-seq peaks were shown to be more consistent predictors of enhancers than H3K4me1/2/3 peaks (46). Therefore, we integrated our ATAC-seq and histone mark ChIP-seq datasets to locate the regulatory elements in the Hydra genome. We report 27,137 candidate regulatory elements, including 3018 candidate enhancer-like elements in the Hydra genome, A subset of the identified regulatory elements are dynamically remodeled during head regeneration. More elements lose accessibility along regeneration time-course. We observed the presence of three open elements near the *Wnt3* locus: a promoter element that gains accessibility during regeneration and two upstream non-promoter open elements. Similarly, 12,983 predicted Hydra genes had at least one non-promoter regulatory element associated with them while 2170 genes had only promoter elements. Thus, 15,153 of the total 33,820 predicted Hydra genes had at least one regulatory element open and detected in our analysis.

Despite the presence of enhancer-like elements in the cnidarian genomes, the CTCF gene is missing in the genomes of both *Nematostella vectensis* (47) and Hydra (using Blast search of the annotated genes). An important future question is to probe the mode of physical interaction of enhancers and promoters in the cnidarian genomes in the absence of CTCF-mediated DNA looping. Additionally, in this study we focused on histone marks associated with active regulatory elements. Future studies on the role of repressive histone marks in gene regulation during important developmental processes, such as head regeneration, can provide further insight.

## Methods and Materials

### RNA-seq Methods and Materials

#### Hydra culture

*Hydra vulgaris* polyps were used for the isolation of RNA. They were fed freshly hatched *Artemia salina* nauplii twice per week and cultured as described previously (48). Animals were starved for at least 1 day before any tissue manipulation or RNA isolation was carried out.

#### Experimental design and tissue manipulation

For each sample, 1-day starved asexual Hydra polyps were selected. For regeneration, 1 animal per sample (with two biological replicates) was bisected at the region 1 (R1) and region 2 (R2) border (Fig 1B) and allowed to undergo head regeneration for a specific period of time (0, 2, 4, 6, 12, 24, or 48 hours). Then the R2-R3 region of the animal of a sample was isolated for RNA extraction. For the budding experiment, the head region of buds from animals at various stages of budding (S1, S3, S4, S5, S6, S7, S8, or S10) (35) was bisected and used for total RNA extraction (Fig 1C). Tissues from tentacles, budding zone, body column, hypostome, and foot were harvested for RNA extraction (Fig 1A).

#### Total RNA extraction

Each isolated tissue was dissolved in Qiagen RNeasy buffer RLT (with 2-betamercaptoethanol added) within 3 minutes of isolation. The dissolved tissue was immediately used for total RNA isolation using Qiagen RNeasy kit according to manufacturer’s protocol. The total RNA for each sample was treated with DNase from TURBO DNA-free kit to remove any genomic DNA contamination. RNA quality was checked with Agilent Bioanalyzer and samples with RIN scores ≥9 were used for RNA-seq library preparation.

#### Illumina library preparation

Multiplexed RNA-seq libraries were built using the Smart-seq2 protocol (49) with slight modifications. Briefly, mRNA from total RNA in each sample was converted to full-length cDNA using poly-dT primer and reverse transcriptase. cDNA was amplified using appropriate number of PCR cycles based on the initial amount of total RNA and as recommended by the Smart-seq2 protocol. 20 ng full-length cDNA for each sample was converted to sequencing library by tagmentation with the Illumina Nextera kit. 8 cycles of PCR were used for library amplification. Libraries were multiplexed and sequenced as 43 bp Illumina paired-end reads.

#### Functional annotation of genes

We used the genome sequence and Augustus predicted gene models from Hydra 2.0 Genome Project (https://research.nhgri.nih.gov/hydra/). The reference transcriptome was annotated with GO terms using Blast2GO (50). First, a BLAST search was done for all the transcripts against NCBI’s non-redundant NR database. The transcripts were then annotated with the GO terms associated with the BLAST hits using the “Mapping” and “Annotation” functions of Blast2GO. The GO terms were expanded using the InterProScan and Annex mapping utilities of Blast2GO.

#### Gene expression analysis

RNA-seq reads for each sample were mapped to the reference transcriptome using Bowtie v. 1.2 (51) with the following parameters: “-X 2000 -a -m 200 -S --seedlen 25 -n 2 -v 3”. Transcript expression levels and read counts were obtained using RSEM v. 1.2.31 (52) with the following parameters: “--paired-end --num-threads 8 --calc-ci”.

Time-series analysis of budding and head regeneration time courses was done using maSigPro v. 1.42.0 (40) and R v. 3.2.3 using the maSigPro functions “p.vector” and “T.fit” with a significance level of 0.01 to find clusters of differentially expressed transcripts and their temporal dynamics.

The heatmap of differentially expressed transcripts was generated using R. The transcripts per millions (TPM) values of the differentially expressed transcripts were log2 transformed and scaled for generating the heatmap.

#### Gene ontology (GO) analysis

Each maSigPro cluster of differentially expressed genes (Fig 3C) was analyzed for GO enrichment using the Fisher’s exact test function of Blast2GO. Each cluster was tested for GO enrichment using the entire reference transcriptome as the reference set. FDR of 5% was used as the significance threshold.

#### Hydra body map differential expression analysis

To identify genes and functions unique to the different body parts of *Hydra*, we did a differential expression analysis and functional annotation of genes upregulated in *Hydra* tentacles, hypostome, body column, budding zone, and foot. We identified differentially expressed genes by doing 10 pairwise comparisons between two different tissues in edgeR (53,54) (Table S1). For each comparison, we filtered for genes having > 1 counts per million per 2 replicates, normalized between samples using TMM normalization (55), and did a false discovery rate correction using the Benjamini-Hochberg procedure. Differentially expressed genes were those which had an FDR > 0.05 and log fold change (logFC) > 2. We merged results of each pairwise comparison to identify genes uniquely upregulated in each of the five tissues (Table S1). Upregulated genes were annotated with potential functions using Blast2GO (50,56,57). Genes were enriched for functions using Fisher’s exact test in Blast2GO using FRD of 5%.

### ATAC-seq and ChIP-seq Methods and Materials

#### Hydra culture

*Hydra vulgaris* polyps were used for the isolation of RNA. They were fed freshly hatched *Artemia salina* nauplii twice per week and cultured as described previously (48). Animals were starved for at least 1 day before any tissue manipulation, nuclei isolation, or crosslinking.

#### Experimental design and tissue manipulation

For chromatin profiling experiments using ATAC-seq, the Hydra polyps were first incubated in a cocktail of four antibiotics for one week with feeding, followed by one week of recovery in sterile medium according to the protocol by Fraune *et al.* (58). This was done to remove commensal bacteria from the Hydra polyps before Tn5 tagmentation during ATAC-seq and avoid contamination from tagmented bacterial DNA.

For each sample, 1-day starved asexual hydra polyps were selected. For regeneration, twenty animals per sample (with two biological replicates) were bisected at the region 1 (R1) and region 2 (R2) border (Fig 1B) and allowed to undergo head regeneration for a specific period of time (0, 2, 4, 6, 12, 24, or 48 hours). Then the R2 regions of the animals of a sample were isolated for nuclei isolation and tagmentation (ATAC-seq) or crosslinking and immunoprecipitation. The following tissues were collected for the body map samples: ten whole animals; foot, budding zone, body column, hypostome, tentacles, and head (hypostome + tentacles) tissues from 20 animals; regenerating R2 tissues from 20 animals were collected for ATAC-seq and ChIP-seq each.

#### Chromatin profiling using ATAC-seq

Nuclei from tissues described in the previous section were isolated using the following protocol based on Endl et al. (59). Briefly, Hydra tissues were washed in ice-cold PBS once. The tissues were homogenized in 1 mL of dissociation medium (3.6 mM KCl, 6 mM CaCl2, 1.2 mM MgSO4, 6 mM sodium citrate, 6 mM sodium pyruvate, 6 mM glucose, 12.5 mM TES, stored in 4°C) in a tissue homogenizer. The solution of homogenized tissue was transferred to an eppendorf tube and centrifuged at 500g for 5 min. The supernatant was removed. The cells were resuspended in 50 μL of cold cell lysis buffer (10 mM Tris-HCl, pH 7.4, 10 mM NaCl, 3 mM MgCl_2_, stored in 4°C) + 0.2% IGEPAL. The sample was immediately spun down at 90g for 8 min. The supernatant was transferred to a fresh eppendorf tube and spun down at 500g for 12 min. The supernatant was removed and the nuclei pellet resuspended in 50 μL ice-cold PBS to remove mitochondrial DNA. The sample was spun down at 500g for 12 min to collect the nuclei for tagmentation. The chromatin in the nuclei pellet was tagmented using 1 μL of Tn5 enzyme and sequencing libraries prepared according to the protocol in Buenrostro et al. (34) with the following modification: DNA fragments in the final sequencing library were size selected for 100-500 bp on a 2% agarose gel. The library qualities were assessed using a Bioanalyzer and the libraries were sequenced as 43-bp paired-end reads on an Illumina NextSeq500. Each sample was performed with two biological replicates.

#### Immunoprecipitation followed by sequencing using ChIP-seq

ChIP-seq libraries were prepared using the ChIPmentation protocol v.1.14 of Schmidl et al. (42). Briefly, tissues described in the section “Experimental design and tissue manipulation” were crosslinked in 1% formaldehyde. The crosslinked chromatin samples were sonicated using A Qsonica sonicator using the following settings: 50%, total 15 minutes, 30 seconds on and 30 seconds off. The chromatin samples were sonicated to an average size range of 200-700 bp. The following antibodies were used for immunoprecipitation: H3K4me3 Rabbit mAb (Cell Signaling Technology, Cat. No. 9751), H3K4me2 Rabbit pAb (Millipore-Sigma, Cat. No. 07030), and H3K27ac Rabbit pAb (Active Motif, Cat. No. 39133). Dynabeads M-280 sheep anti-rabbit IgG beads (Thermo Fisher Scientific, Cat. No. 11203D) were used for immunoprecipitation. The sequencing libraries were prepared according to protocol. The library qualities were on a Bioanalyzer and the libraries were sequenced as 43-bp paired-end reads on an Illumina NextSeq500. Each experiment was performed with two biological replicates.

#### ATAC-seq data analysis

Adapter sequences and low quality base pairs from the paired-ends reads were trimmed using Trimmomatic v. 0.35 (60) using the following parameters: “PE [read1.fastq] [read2.fastq] pe_read1.fastq.gz se_read1.fastq.gz pe_read2.fastq.gz se_read2.fastq.gz ILLUMINACLIP:NexteraPE-PE.fa:2:30:8:4:true LEADING:20 TRAILING:20 SLIDINGWINDOW:4:17 MINLEN:30”. The trimmed reads were first mapped to Hydra mitochondrial DNA sequences to filter the mitochondrial reads. The unmapped reads were mapped to the genome sequence from Hydra 2.0 Genome Project (https://research.nhgri.nih.gov/hydra/) using Bowtie v1.2 (51) with the following parameters: -X 2000 -v 3 -m 3 -k 1 –best. Peaks were called using Homer (41) using the following parameters: - size 500 -minDist 50 -fdr 0.01 -style factor. Overlapping peaks between the two replicates of each sample were kept for downstream analysis. The peaks from all samples were merged using Bedtools v.2.23.0 (61). The read coverage of peaks for each sample was obtained using “coverageBed” function of Bedtools. The read counts for each sample were normalized for efficiency (number of reads within peaks divided by total number of mapped reads) and reads per million. Differentially accessible (DA) peaks were determined using edgeR’s GLM function (53) at the significance level of 5% FDR and minimum 2-fold change. Signal densities at the peaks were plotted using Deeptools (62). Hydra genes were associated with the ATAC-seq peaks using “annotatePeaks.pl” from Homer package.

#### ChIP-seq data analysis

Adapter sequences and low quality base pairs from the paired-ends reads were trimmed using Trimmomatic v. 0.35 (60) using the parameters in the previous section. The trimmed reads were mapped to the genome sequence from Hydra 2.0 Genome Project (https://research.nhgri.nih.gov/hydra/) using Bowtie v. 1.2 (51) with the following parameters: - X 2000 -v 3 -m 3 -k 1 –best. The read coverage of ATAC-seq peaks for each sample was obtained using “coverageBed” function of Bedtools. The read counts for each sample were normalized for efficiency (number of reads within peaks divided by total number of mapped reads) and reads per million. Signal densities of the histone marks at the ATAC-seq peaks were plotted using Deeptools (62).

---

**Figure S1:**
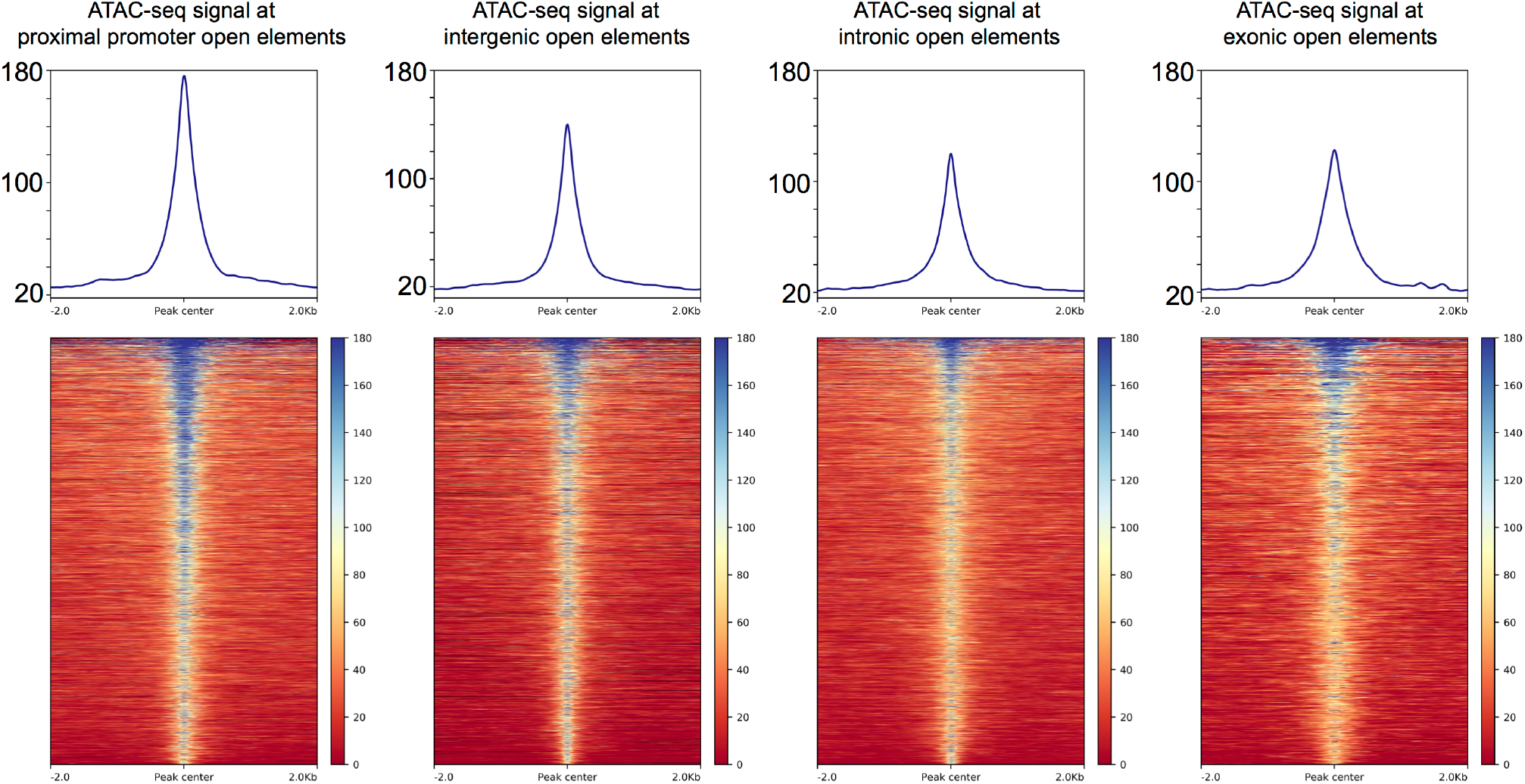
ATAC-seq signal enrichment at each type of open-chromatin element. All graphs and heatmaps are to the same scale.

